# EEG and behavioral correlates of attentional processing while walking and navigating naturalistic environments

**DOI:** 10.1101/2021.05.27.445993

**Authors:** Magnus Liebherr, Andrew W. Corcoran, Phillip M. Alday, Scott Coussens, Valeria Bellan, Caitlin A. Howlett, Maarten A. Immink, Mark Kohler, Matthias Schlesewsky, Ina Bornkessel-Schlesewsky

**Affiliations:** Department of Neuroscience, Karolinska Institutet, Stockholm, Sweden; Department of General Psychology: Cognition, University Duisburg-Essen, Germany; Cognitive and Systems Neuroscience Research Hub, University of South Australia, Adelaide, Australia; Cognition and Philosophy Laboratory, Monash University, Melbourne, Australia; Innovation, Implementation & Clinical Translation (IIMPACT) in Health, University of South Australia, Adelaide, Australia; Sport, Health, Activity, Performance and Exercise Research Centre, Flinders University, Adelaide, Australia; School of Psychology, University of Adelaide, Adelaide, Australia

**Keywords:** Mobile EEG, ERP, attention, real-world, cognitive load

## Abstract

The capacity to regulate one’s attention in accordance with fluctuating task demands and environmental contexts is an essential feature of adaptive behavior. Although the electrophysiological correlates of attentional processing have been extensively studied in the laboratory, relatively little is known about the way they unfold under more variable, ecologically-valid conditions. Accordingly, this study employed a ‘real-world’ EEG design to investigate how attentional processing varies under increasing cognitive, motor, and environmental demands. Forty-four participants were exposed to an auditory oddball task while (1) sitting in a quiet room inside the lab, (2) walking around a sports field, and (3) wayfinding across a university campus. In each condition, participants were instructed to either count or ignore oddball stimuli. While behavioral performance was similar across the lab and field conditions, oddball count accuracy was significantly reduced in the campus condition. Moreover, event-related potential components (mismatch negativity and P3) elicited in both ‘real-world’ settings differed significantly from those obtained under laboratory conditions. These findings demonstrate the impact of environmental factors on attentional processing during simultaneously-performed motor and cognitive tasks, highlighting the value of incorporating dynamic and unpredictable contexts within naturalistic designs.

## Introduction

Attention is a finite resource that must be strategically managed throughout daily life. The hyper-connected nature of the modern world, and in particular, the ubiquitous devices, apps, and media that constantly vie for our attention, expose us to an abundance of information that must be selectively sampled, navigated, and at times, held in abeyance. Arguably, it has never been more important to understand the impact of competing attentional demands on cognitive function and motor action. The present study builds on recent developments in mobile electroencephalography (EEG) by investigating the psychophysiological effects of cognitive task load, motor demands, and environmental complexity on attention in real-world scenarios.

A great deal of knowledge about the nature of attention and the influence of distracting stimuli has been derived from carefully controlled laboratory studies ^1^. By contrast, much less is known about the characteristics of attention under real-world situations. However, recent technological developments in neurophysiological measures (e.g., EEG and eye-tracking) have enabled researchers to address such topics outside the laboratory (e.g., while driving) ^2^. The present study exploits this technical progress to examine how event-related potentials (ERPs) relating to sensory and attentional processing differ across laboratory and more ecologically-valid situations. Here, we focus on the P3 component, which has been most frequently investigated in previous real-world studies ^3^, and the mismatch negativity (MMN), which constitutes an index of early sensory processing ^4^.

The P3 constitutes the third positive deflection after stimulus presentation, typically occurring at a latency ranging 250 to 500 ms and most dominantly in centro-parietal areas ^5,6^. This component has been related to updating and categorization processes ^7^, the intensity of performance ^8,9^, as well as the amount of resources dedicated to task performance ^10,11^. Furthermore, the P3 can serve as an index of the distribution of attention in competing concurrent tasks ^12^ and has been shown to be sensitive to alterations in task difficulty ^13^.

The MMN is elicited in the presence of any discriminable change in some repetitive aspect of stimulation. Peaking within 100 to 250 ms from change onset, the MMN exhibits the strongest intensity in temporal and frontal areas of topographic scalp maps, and can be detected even when the participant pays no attention to the task ^14–16^. Although the MMN has been most intensively studied in the auditory modality ^17^, previous studies have also identified the component in visual settings ^18^.

Traditionally, the P3 and MMN components are most frequently studied using the oddball paradigm. Within this paradigm, participants are typically presented with an infrequently occurring target stimulus (e.g., a high-pitch tone) embedded within sequences of frequently presented distractor stimuli (e.g., low-pitch tones). While both the P3 and the MMN are elicited by such target stimuli, the MMN can also be detected when the individual is not actively attending to the tone sequence ^19^.

From pioneers to contemporary studies, the oddball paradigm has also been frequently used to investigate the influence of motor demands (e.g., varying gait parameters) on ERPs, where participants are asked to perform the task while walking on a treadmill with varying velocity (slow walking vs. fast walking) ^20^. Here, findings suggest that the amplitude, latency and topography of event-related potentials (N1, N2, P3) vary depending on the difficulty of both the motor and the cognitive task ^20,21^. When occurring in parallel with a cognitive task, motor demands (e.g., during walking) can thus be viewed as inducing a dual-task setting that calls for appropriate allocation of attention between cognitive and motor systems ^22^.

In recent years, studies have begun to measure ERPs during dual-task scenarios performed under more naturalistic conditions, such as cycling ^23,24^, skateboarding ^25^, or walking ^26^ outside the lab (e.g., on empty corridors, parking places, or the lawn on the outside premises). Such findings demonstrate a relation between attentional processing and the cognitive load induced by each situation. For example, cognitive task performance, parietal P3 amplitude, and frontal midline theta power have been found to decrease with progressively increased situational load (i.e., standing vs. walking on a treadmill vs. walking along a hallway) ^27^. Furthermore, these findings show a greater impact of environmental factors on electrophysiological markers of attention compared to motor demands (i.e., being wheeled along a hallway vs. walking on a treadmill). To date, however, most real-world studies on walking involve fairly undemanding environmental settings, such as an empty straight corridor ^27^ or the lawn of an institute’s grounds ^3^.

Within a more naturalistic context, Aspinall and colleagues described EEG changes in terms of increased engagement and arousal in more busy environmental settings (i.e., urban shopping street and busy commercial district) compared to green spaces ^28^. However, the interpretation of these findings is complicated by the authors’ reliance on proprietary EEG metrics that are not commonly reported in the literature. More recently, Pizzamiglio and colleagues reported increased theta and beta power when participants were walking across a campus while conversing, as compared to standing ^29^. However, studies that have analyzed attention-related ERP responses under such ecologically valid conditions are rare and often limited to virtual reality ^22^.

In sum, previous studies on walking in naturalistic environments have either focused on the effect of motor demands ^3^, used relatively monotonous scenarios (e.g., empty corridors) ^27^, or did not analyze attention-related ERPs ^29–31^. Here, we sought to fill an important gap in this literature by investigating the effects of motor demands, environmental complexity, and cognitive task load on such ERPs. We report findings from an auditory oddball paradigm in which participants either attended to or ignored experimental stimuli while engaging in three situational contexts: (1) ordinary laboratory conditions, (2) walking around a sports field, and (3) navigating a route through a university campus. Behavioral data and ERPs pertaining to the MMN and P3 were analyzed to investigate the effect of each experimental manipulation on task performance and attentional processing.

Based on previous findings from traditional laboratory studies and studies in increasingly naturalistic environments, we hypothesized a decrease in P3 and MMN amplitude as well as task-performance with increased environmental and motor complexity, due to the reduction in attentional resources available for cognitive task performance. In addition, we predicted that P3 and MMN effects would be more pronounced under conditions in which specific emphasis is placed on the cognitive task, compared to when the tone stimuli are ignored.

## Methods

### Participants

Healthy, right-handed, native English-speaking adults aged 18-40 years were eligible to participate in this study. Of the 44 recruited participants, one was excluded due to left-handedness, and a further two were excluded due to missing EEG data. The remaining sample consisted of 26 females and 15 males aged 18–39 years (*M* = 23.1 years, *SD* = 4.8). All participants reported normal (or corrected-to-normal) vision and audition, no history of neurological or psychological disorders, and no diagnosed intellectual or language disorder.

The study was performed in accordance with the ethical standards laid down in the Declaration of Helsinki and approved by the local ethics committee at the University of South Australia (ID: 0000034991). All participants provided written informed consent prior to the experiment and were informed that they could end participation at any time without reprisal.

### Materials

The auditory oddball task comprised two distinct tones, one presented frequently (standard) and the other infrequently (deviant). These stimuli were a French horn tone (Dominant frequency of 1046.502Hz, High C or C6) and a piano tone (Dominant frequency of 523.251Hz, Middle C or C5). We used complex tones because they constitute more naturalistic stimuli than pure tones ^32^.

Tones were presented binaurally via conventional in-ear headphones (Anko) for 150 ms, with an interstimulus interval of 350 ms. One-hundred and fifty stimuli were presented per block, of which 13-30% were deviants (*M* = 20%). Tones were randomly assigned as the standard/deviant. Stimulus presentation was driven by OpenSesame v3.0.7 ^33^ running on a commercial compact laptop in conjunction with BrainVision StimTrak (Gain = 10, Trigger Level = 1V). The volume of the oddball stimuli was kept the same for all participants and conditions (approximately 60 dB).

### Procedure

After providing written, informed consent, participants completed a demographic questionnaire and the Edinburgh Handedness Inventory ^34^, followed by the EEG setup. Before starting the task, the two tones were presented to familiarize participants with the stimuli. Participants performed 5 blocks of the oddball task in each of the three environments (Lab, Field, Campus) while counting the number of deviant stimuli (Count), and again in each of the three environments while ignoring task stimuli (Ignore). The order of the task manipulation was randomized, as was the order of environmental conditions nested within either task condition. We chose counting rather than button press responses as our measure of attentional engagement because previous studies have reported that P3 topography is confounded by motor-evoked potentials ^35^.

Within the Lab condition, participants performed the oddball task while sitting alone in an interview room. The room was sparsely furnished with a large table, six chairs and a sideboard, surrounded by three white walls without pictures, and one frosted glass wall. Participants sat on a chair with armrests, with the laptop in front of them (supported by a harness strapped to their torso). The participant’s gaze was directed towards a white wall. The glass wall, which separated the interview room from a corridor, was on their right.

The Field condition involved walking around a grass sports field at the Magill campus of the University of South Australia, Adelaide. The sports field is surrounded by a grandstand, a car park for members of the university, trees and fencing (see Figure 1a). The Campus condition involved navigating a pre-defined route through the university campus adjacent to the sports field. Participants were provided a map with a cross-marked destination (red point) to which they should navigate via several waypoints (yellow points) (see Figure 1b). The route included both outdoor and indoor sections. Participants had to use stairs, open doors, and interact with other people on the campus.

**Figure 1.**
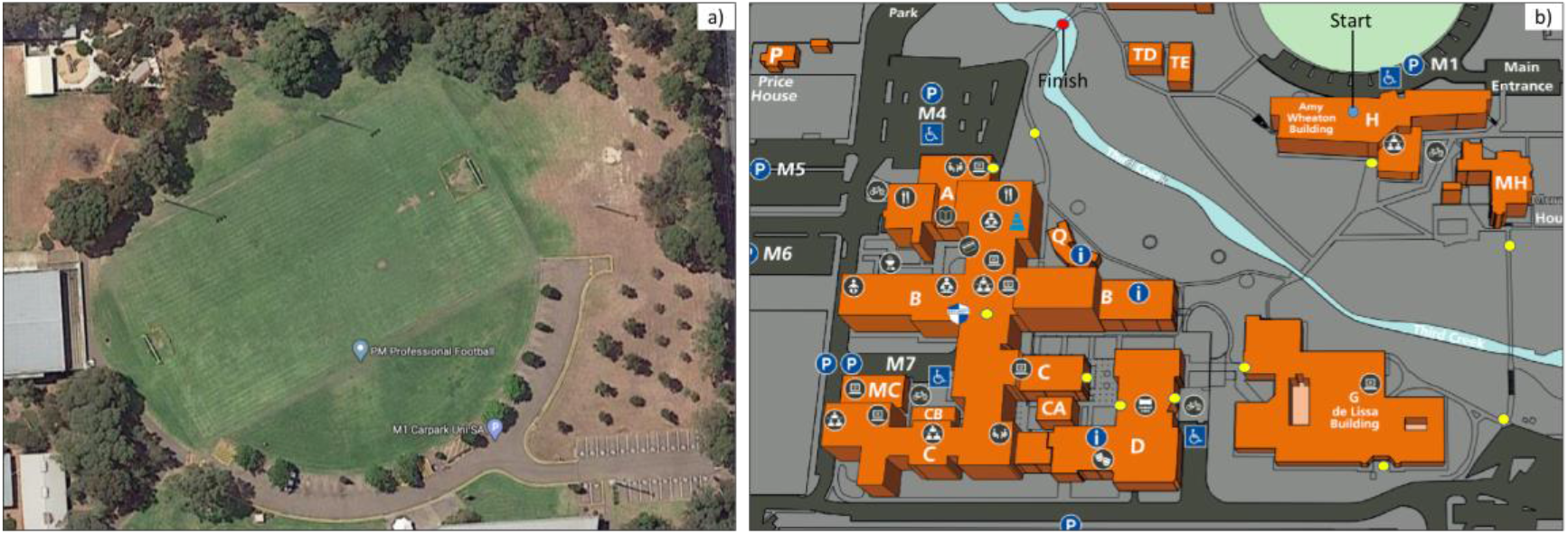
a) Grass sports field at the Magill campus at the University of South Australia (from *Google Maps*, by Google, https://bit.ly/2R0XXjl), b) Reproduction of the map provided to the participants in the “campus condition”, showing the route across the Magill campus. Participants began at the blue dot that represents the starting point and had to follow the yellow dots until they reached the red dot that represents the finish point.

During both walking conditions, participants carried the laptop in front of them (supported by the harness) while carrying the StimTrak in a plastic pocket (attached with a lanyard) over one shoulder as they made their way along the route. Participants were permitted to walk at a comfortable, self-determined pace. The entire testing session lasted between 2.5 and 3 hours, with sufficient breaks between conditions to avoid fatigue. Data collection occurred during the summer months, with temperatures typically ranging between 25 and 30°C. Sessions were not performed when it was raining or in extreme temperatures.

### EEG acquisition and preprocessing

The EEG was continuously recorded from 32 Ag/AgCl actiCAP mounted active electrodes using a Brain Vision LiveAmp 32 amplifier (Brain Products GmbH, Gilching, Germany). Electrode positions conformed to the extended 10–20 system: Fp1, Fz, F3, F7, FT9, FC5, FC1, C3, T7, TP9, CP5, CP1, Pz, P3, P7, O1, Oz, O2, P4, P8, TP10, CP6, CP2, Cz, C4, T8, FT10, FC6, FC2, F4, F8, Fp2. All channels were digitized at a sampling rate of 500 Hz. AFz served as the ground and FCz as the online reference. Electrode impedances were below 20 kΩ at the beginning of the experiment. EEG data were stored on a micro SD card inside the amplifier; the amplifier was kept inside a mesh pocket on the back of the EEG cap during the experiment. After the experiment, the data were transferred from the SD card to a hard drive for offline processing and analysis.

EEG data were preprocessed in MATLAB R2019b (v9.7.0.1319299; The MathWorks, Natick, MA) using custom-built scripts incorporating functions from the EEGLAB (v2019.1;^36^) and ERPLAB (v8.10;^37^) toolboxes. Marker files were first manually edited to remove artefactual triggers caused by feedback from the auditory equipment. Edited data were then imported into EEGLAB and downsampled to 250 Hz. Accelerometer channels and non-experimental recording segments were excluded. The online reference was then reintroduced into the dataset, and the data re-referenced to the average of TP9 and TP10 in order to approximate linked mastoids. The data were then high-pass filtered (passband edge = 0.5 Hz, transition width = 0.5 Hz) using a zero-phase, hamming-windowed, finite impulse response (FIR) sinc filter (implemented in the *pop_eegfiltnew* function from the *firfilt* plugin; v2.4).

Filtered data were processed with artifact subspace reconstruction (implemented with the *pop_clean_artifacts* function from the *clean_rawdata* plugin; v2.3). This routine was used to remove flat or noisy channels (flatline criterion = 5 s; channel correlation criterion = 0.80; line noise criterion = 4), and to correct for large amplitude artifacts (burst criterion = 10, ^38,39^). Data were then low-pass FIR filtered (passband edge = 40 Hz, transition width = 10 Hz) prior to independent component analysis (*binica* implementation of the extended logistic infomax ICA algorithm,^40,41^). Resultant decompositions were automatically classified using the ICLabel plugin (v1.3, ^42^). Components classed as either ocular or cardiac in origin with >90% probability were subtracted (median = 2, range = 1-3).

Missing channels (median = 1, range = 0-4) were interpolated for the purposes of ERP visualization but excluded from statistical analysis. All EEG event markers were systematically shifted -200 ms relative to their recorded latencies (*pop_erplabShiftEventCodes*) to account for the lag in signal transmission introduced by the StimTrak. The EEG was then epoched [-200, 800 ms] relative to the corrected event markers. Extracted epochs were mean-centered and subjected to a moving window artefact rejection routine (*pop_artmwppth*; voltage threshold = 125 µV, time period = [-200, 600 ms]; window size = 100 ms, window step = 50 ms). Grand average ERP waveforms and difference waves, along with their accompanying standard error estimates, were computed using the *pop_gaverager* function (weighted averages based on number of available trials per participant). Finally, mean voltages were extracted from three time-windows per trial epoch (Prestimulus = [-200, 0 ms]; MMN = [100, 250 ms]; P3 = [250, 500 ms]) and exported for statistical analysis.

### Data analysis

All statistical analysis was conducted in *R* (v3.6.2 ^43^) using the *RStudio Desktop IDE* (v1.2.5033 ^44^). Behavioral and EEG data were analyzed using linear mixed-effects models fit using the *lmer()* function from *lme4* (v1.1-23 ^45^). All mixed-effects models included a by-participant random intercept term to account for non-independence of (within-subject) observations. All categorical independent variables were entered as unordered factors using sum-to-zero contrast coding (reference category coded -1).

For the behavioral model, deviant counts for each block of the Count task were included as the dependent variable, while number of deviant stimuli and environmental condition were included as independent variables. Note that including the number of presented deviants as a covariate controlled for differences in the ground truth count across blocks, enabling us to implicitly model count accuracy as a function of environmental condition.

Separate EEG models were fitted for the MMN and P3 time-windows, with single-trial mean voltage estimates serving as the dependent variable. Environmental condition, stimulus type, and task manipulation were entered as independent variables and permitted to interact with one another. Models additionally included mean prestimulus voltage as a scaled covariate in order to address trial-by-trial variation in prestimulus activity (essentially functioning as a form of baseline correction; see ^46^), and by-channel random intercepts to account for topographic differences in voltage estimates (see supplementary materials for models that specify midline channel location as a fixed effect). By-participant random slopes of environment were included in the MMN model; by-participant random slopes of environment and stimulus were included in the P3 model (more complex random effects structures failed to converge).

Main effects and interaction terms were assessed using Wald tests from Type-II ANOVA tables obtained from the *car* package (v3.0-10 ^47^) Model summary tables were produced using *sjPlot* (v2.8.7 ^48^). P-values for model terms were derived using Satterthwaite’s method for approximating degrees of freedom. Post-hoc contrasts were obtained via the *emmeans* package (v1.5.1 ^49^). Pairwise contrasts were corrected using the Tukey method; interaction contrasts were corrected using the Sidak method. Plotted model predictions are estimated marginal means (also obtained from the *emmeans* package) and visualised with the aid of *tidyverse* (v1.3.0 ^50^). Model predictions are plotted with 84% confidence intervals, where non-overlapping intervals indicate a significant difference at α = .05 ^51^.

### Data Availability Statement

Raw and preprocessed datasets, along with all code necessary to reproduce the analysis pipeline reported in this paper, are openly available from the OpenNeuro database [Dataset: doi: 10.18112/openneuro.ds003620.v1.0.2]. Data were archived in accordance with the Brain Imaging Data Structure (BIDS) standard for EEG ^52^ with the assistance of functions from the bids-matlab-tools plugin (v5.2; https://github.com/sccn/bids-matlab-tools).

## Results

### Behavioral data

Two participants were excluded from analysis due to numerous highly inaccurate count estimates (more than double the number of presented deviants on the majority of blocks). Due to a programming error, count estimates from the final block of trials were missing for 21 participants. A further 16 count estimates were excluded as outliers (> 3 *SD* from the sample mean after estimates had been normalized by the number of presented deviants). The included 39 participants contributed a median 14 counts to the model.

Numerically, mean count estimates were similar across Lab and Field conditions, but reduced in the Campus condition. Linear mixed-effects modelling revealed a significant effect of Environment (χ^2^(2)=12.08, *p*=.002) after controlling for the number of presented deviants. Post-hoc contrasts revealed that count estimates were significantly reduced during the Campus condition compared to both the Lab (*p*_adj_ =.005) and Field (*p*_adj_ =.014) conditions. No significant difference was found between Lab and Field conditions (*p*_adj_ =.930).

### Event related potentials

A further three participants were excluded from the ERP analysis due to excessively noisy recordings (>50% epochs rejected due to artifact). Epoch rejection rates for the remaining 36 participants ranged from 0 to 38.6% (*M* = 7.2, *SD* = 9.9).

Grand average ERPs and difference waveforms from electrodes Fz and Cz are displayed in Figure 3 (see supplementary materials for additional ERP plots). In the Lab condition, both channels revealed a marked negative-going waveform in the early time-window for deviant stimuli, consistent with a MMN component. This effect was evident irrespective of task manipulation, although scalp topographies displaying the mean voltage difference between stimuli indicate that the effect was more pronounced over centro-parietal sensors during the Count task (see Figure 3). Conversely, deviants elicited a strong positive deflection in the first half of the later time-window, consistent with a P3 response. This effect was also strongest over centro-parietal channels in the Count task.

**Figure 2.**
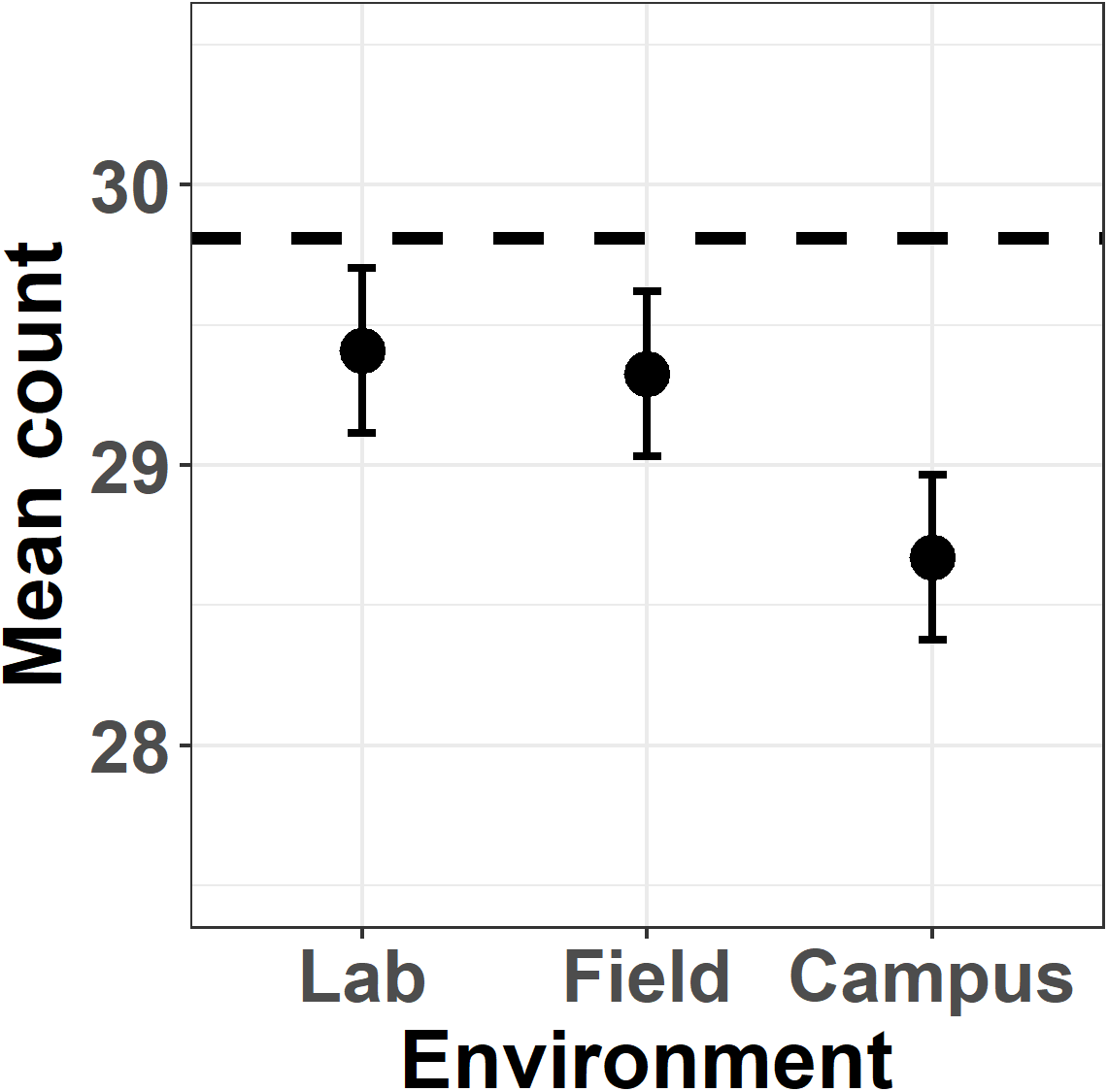
Estimated marginal mean counts of deviant stimuli per block across environmental conditions (Lab, Field, Campus). Error bars indicate 84% confidence intervals. Dashed line indicates perfect performance.

**Figure 3.**
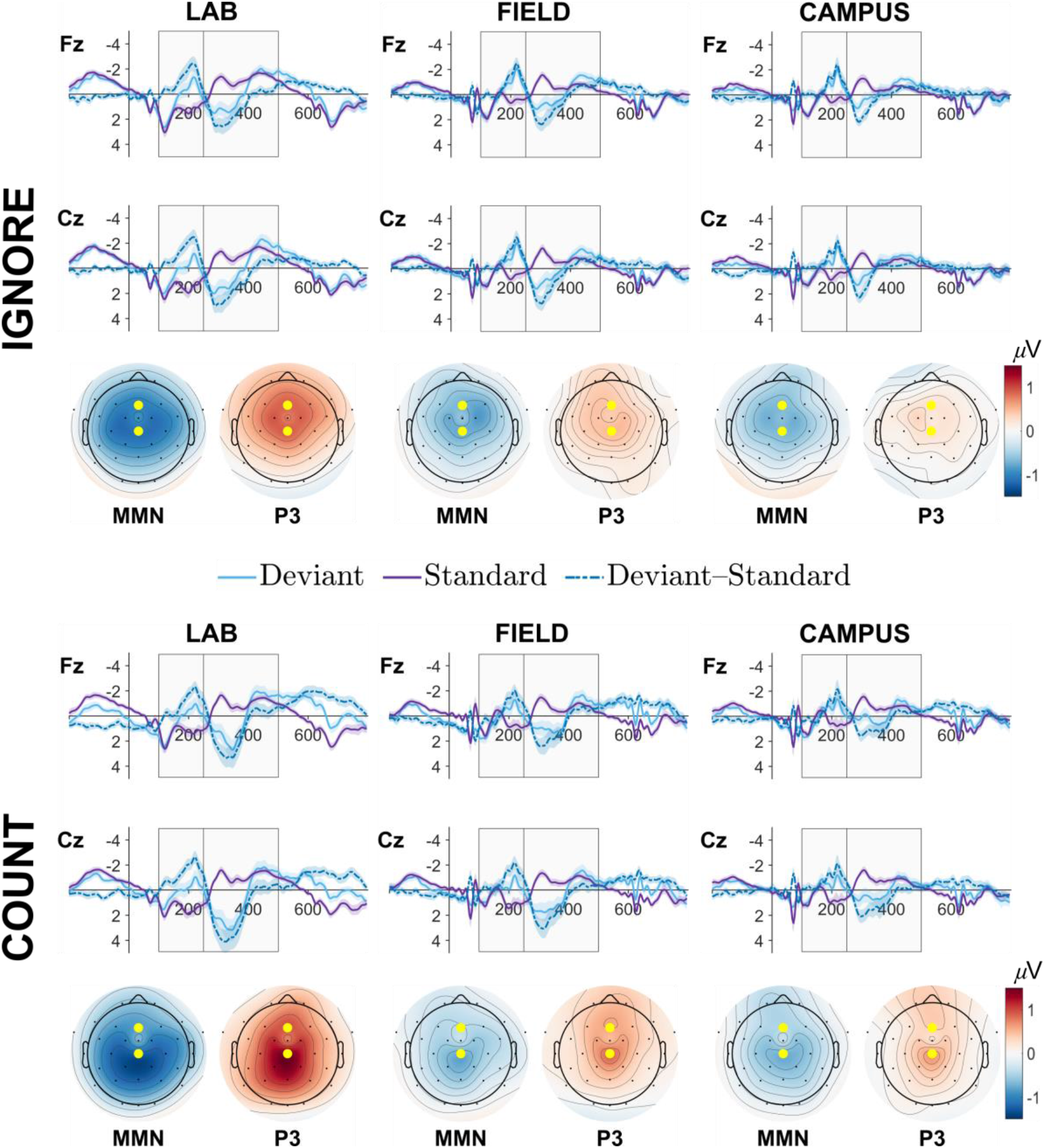
Grand average event-related potentials (ERPs; solid lines), difference waveforms (broken lines) and topographies for Ignore (top panel) and Count (bottom panel) tasks across Lab (left column), Field (middle column), and Campus (right column) conditions. Light blue waveforms depict ERPs evoked by deviant stimuli; purple waveforms depict ERPs evoked by standard stimuli. Broken lines indicate the difference in mean voltage (*µ*V) between deviants and standards. Topographies depict mean voltage differences (*µ*V) averaged across the two time-windows of interest (MMN = 100 – 250 ms; P3 = 250 – 500 ms). Yellow dots indicate electrode locations for plotted ERPs (upper = Fz, lower = Cz).

Field and Campus conditions rendered qualitatively similar ERP waveforms. Deflections in the MMN time-window were positive-going for standard tones, and negative-going for deviants, rendering a negative difference wave that was most pronounced over central electrode sites. This pattern of activity was again reversed in the P3 time-window, resulting in a centrally distributed positivity that peaked approximately 300 ms post stimulus onset. This effect was less pronounced in the Campus compared to the Field condition. Comparing across task manipulations, the difference between deviant and standard waveforms in the MMN time-window was slightly accentuated, and more frontally distributed, in the Ignore task, while the P3 window revealed slight accentuation over centro-parietal regions in the Count task.

Taken together, the qualitative pattern of both the early and late ERP components elicited during the walking conditions was generally similar to those elicited in the Lab condition, but of reduced magnitude.

### MMN time-window

Type II Wald tests revealed a significant three-way interaction between stimulus, task, and environmental condition for the MMN model (χ^2^ (2) = 7.67, *p* = .022; see Table 1 for full decomposition). This interaction is resolved in Figure 4 (see Table 2 for the full model summary). This plot reveals that mean voltage was significantly more negative for all pairwise contrasts of stimulus type within each factorial combination of Task and Environment (all *p*_adj_ <.001), consistent with the elicitation of the MMN. In line with the ERP waveforms depicted in Figure 3, the magnitude of this difference was greatest in the Lab condition, and reduced in the other two conditions. This diminution was predominantly driven by the modulation of cortical responses to standard stimuli, which were less positive-going in the Field and Campus conditions compared to the corresponding Lab condition, irrespective of task instructions (all *p*_adj_ <.001).

**Table 1.** Analysis of deviance tables for linear mixed-effects models evaluating voltage changes in the MMN (100-250 ms) and P3 (250-500 ms) time-windows.

|  | MMN time-window |  |  | P3 time-window |  |  |
| --- | --- | --- | --- | --- | --- | --- |
| | $\chi^2$ | *d.f.* | *p-value* | $\chi^2$ | *d.f.* | *p-value* |
| stimulus | 5176.37 | 1 | <.001 | 36.39 | 1 | <.001 |
| task | 0.87 | 1 | .350 | 76.78 | 1 | <.001 |
| environ | 27.66 | 2 | <.001 | 8.73 | 2 | .013 |
| prestim.s | 246102.39 | 1 | <.001 | 575055.15 | 1 | <.001 |
| stimulus:task | 0.07 | 1 | .786 | 221.33 | 1 | <.001 |
| stimulus:environ | 799.65 | 2 | <.001 | 1775.21 | 2 | <.001 |
| task:environ | 105.34 | 2 | <.001 | 492.80 | 2 | <.001 |
| stimulus:task:<br>environ | 7.67 | 2 | .022 | 33.40 | 2 | <.001 |

Note. $\chi^2$ = Type II Wald $\chi^2$ test statistic, *d.f.* = degrees of freedom, *stimulus* = oddball task stimulus (Deviant, Standard), *task* = cognitive task condition (Count, Ignore), *environ* = environmental context (Lab, Field, Campus), *prestim.s* = prestimulus EEG voltage (scaled).

**Table 2.**
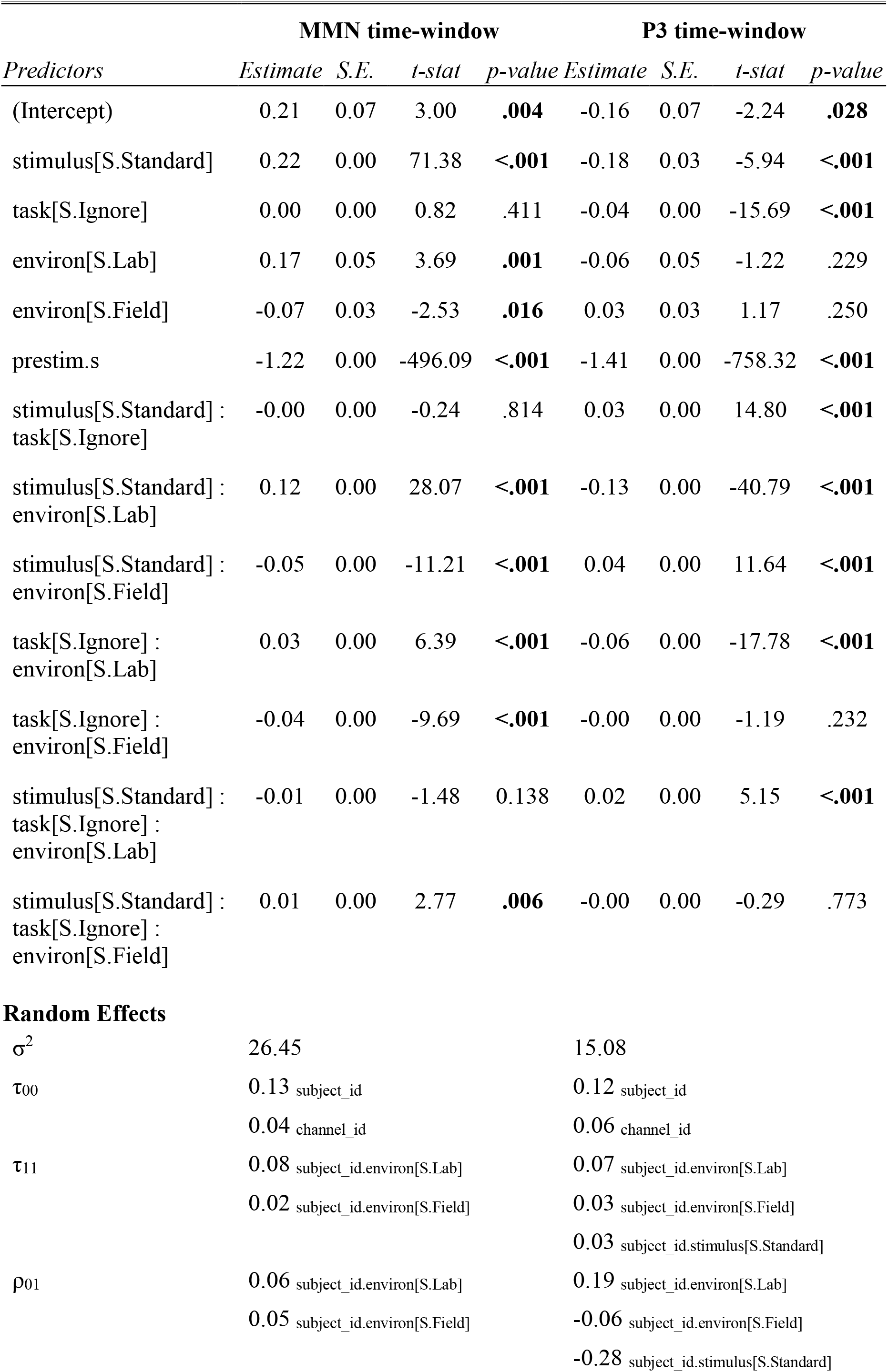
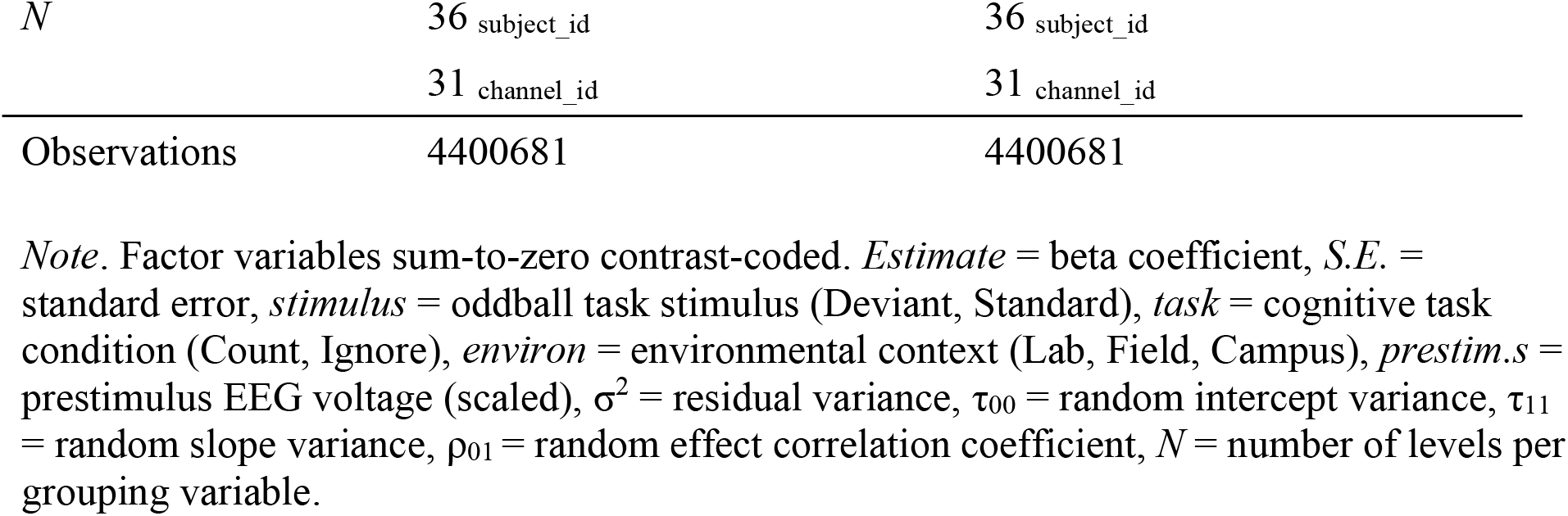
Summary of linear mixed-effects models for the MMN and P3 time-windows.

| *Predictors* | **MMN time-window** |  |  |  | **P3 time-window** |  |  |  |
| --- | --- | --- | --- | --- | --- | --- | --- | --- |
|  | *Estimate* | *S.E.* | *t-stat* | *p-value* | *Estimate* | *S.E.* | *t-stat* | *p-value* |
| (Intercept) | 0.21 | 0.07 | 3.00 | **.004** | -0.16 | 0.07 | -2.24 | **.028** |
| stimulus[S.Standard] | 0.22 | 0.00 | 71.38 | **<.001** | -0.18 | 0.03 | -5.94 | **<.001** |
| task[S.Ignore] | 0.00 | 0.00 | 0.82 | .411 | -0.04 | 0.00 | -15.69 | **<.001** |
| environ[S.Lab] | 0.17 | 0.05 | 3.69 | **.001** | -0.06 | 0.05 | -1.22 | .229 |
| environ[S.Field] | -0.07 | 0.03 | -2.53 | **.016** | 0.03 | 0.03 | 1.17 | .250 |
| prestim.s | -1.22 | 0.00 | -496.09 | **<.001** | -1.41 | 0.00 | -758.32 | **<.001** |
| stimulus[S.Standard] :<br>task[S.Ignore] | -0.00 | 0.00 | -0.24 | .814 | 0.03 | 0.00 | 14.80 | **<.001** |
| stimulus[S.Standard] :<br>environ[S.Lab] | 0.12 | 0.00 | 28.07 | **<.001** | -0.13 | 0.00 | -40.79 | **<.001** |
| stimulus[S.Standard] :<br>environ[S.Field] | -0.05 | 0.00 | -11.21 | **<.001** | 0.04 | 0.00 | 11.64 | **<.001** |
| task[S.Ignore] :<br>environ[S.Lab] | 0.03 | 0.00 | 6.39 | **<.001** | -0.06 | 0.00 | -17.78 | **<.001** |
| task[S.Ignore] :<br>environ[S.Field] | -0.04 | 0.00 | -9.69 | **<.001** | -0.00 | 0.00 | -1.19 | .232 |
| stimulus[S.Standard] :<br>task[S.Ignore] :<br>environ[S.Lab] | -0.01 | 0.00 | -1.48 | 0.138 | 0.02 | 0.00 | 5.15 | **<.001** |
| stimulus[S.Standard] :<br>task[S.Ignore] :<br>environ[S.Field] | 0.01 | 0.00 | 2.77 | **.006** | -0.00 | 0.00 | -0.29 | .773 |
| **Random Effects** |  |  |  |  |  |  |  |  |
| $\sigma^2$ | 26.45 | | | | 15.08 | | | |
| $\tau_{00}$ | 0.13 subject_id | | | | 0.12 subject_id | | | |
|  | 0.04 channel_id |  |  |  | 0.06 channel_id |  |  |  |
| $\tau_{11}$ | 0.08 subject_id.environ[S.Lab] | | | | 0.07 subject_id.environ[S.Lab] | | | |
|  | 0.02 subject_id.environ[S.Field] |  |  |  | 0.03 subject_id.environ[S.Field] |  |  |  |
|  |  |  |  |  | 0.03 subject_id.stimulus[S.Standard] |  |  |  |
| $\rho_{01}$ | 0.06 subject_id.environ[S.Lab] | | | | 0.19 subject_id.environ[S.Lab] | | | |
|  | 0.05 subject_id.environ[S.Field] |  |  |  | -0.06 subject_id.environ[S.Field] |  |  |  |
|  |  |  |  |  | -0.28 subject_id.stimulus[S.Standard] |  |  |  |

|  |  |  |
| --- | --- | --- |
| $N$ | 36 subject_id | 36 subject_id |
|  | 31 channel_id | 31 channel_id |
| Observations | 4400681 | 4400681 |

*Note.* Factor variables sum-to-zero contrast-coded. *Estimate* = beta coefficient, *S.E.* = standard error, *stimulus* = oddball task stimulus (Deviant, Standard), *task* = cognitive task condition (Count, Ignore), *environ* = environmental context (Lab, Field, Campus), *prestim.s* = prestimulus EEG voltage (scaled), $\sigma^2$ = residual variance, $\tau_{00}$ = random intercept variance, $\tau_{11}$ = random slope variance, $\rho_{01}$ = random effect correlation coefficient, $N$ = number of levels per grouping variable.

**Figure 4.**
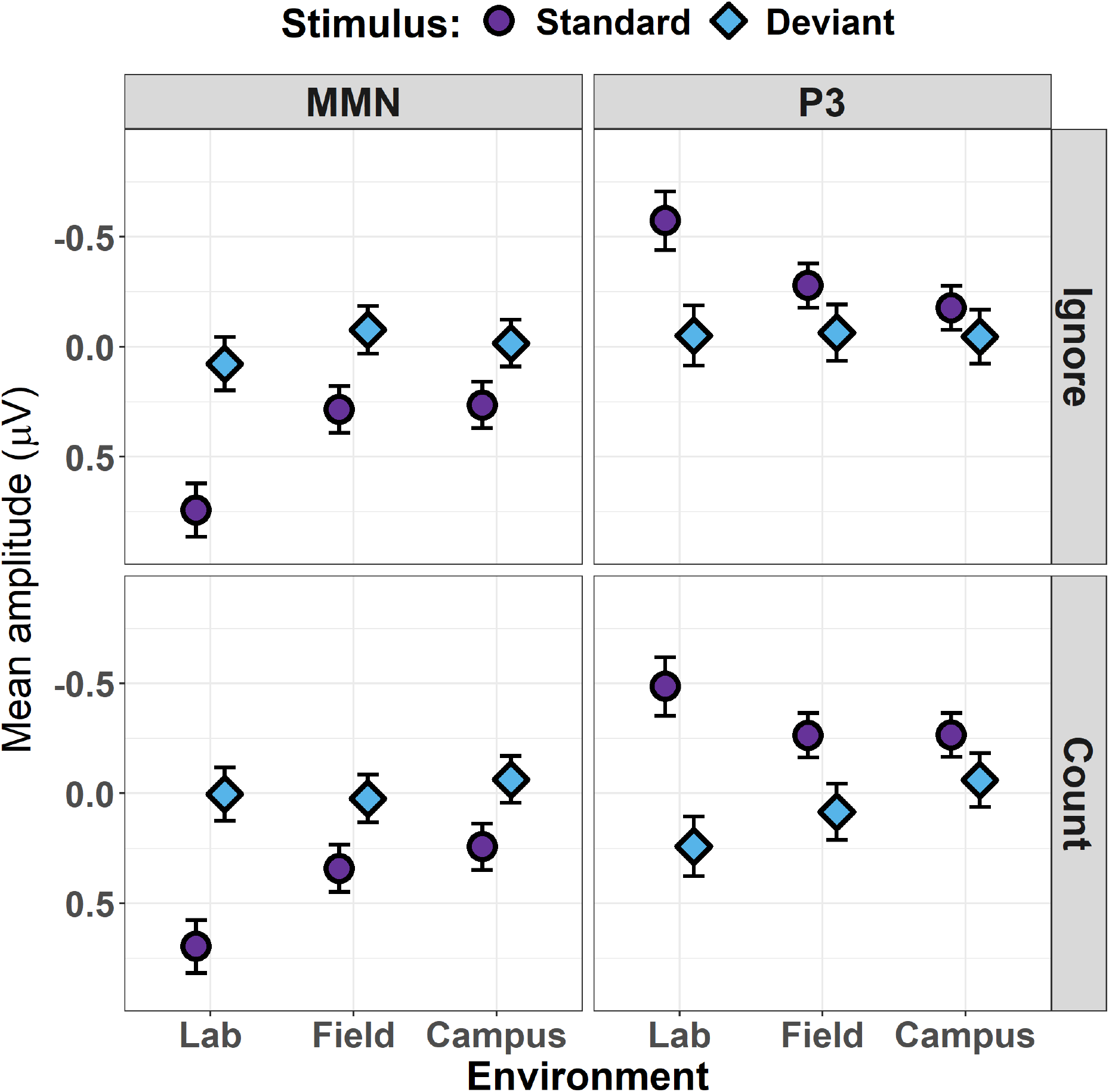
Estimated marginal mean voltage amplitudes for MMN (left column) and P3 (right column) time-windows during the Ignore (top row) and Count (bottom row) conditions of the oddball task. Responses to standard tone stimuli are depicted by purple circles; responses to deviant tone stimuli are depicted by blue diamonds. Estimates for each environmental condition (Lab, Field, Campus) are displayed along the x-axis of each subplot. Error bars indicate 84% confidence intervals.

The difference between stimulus responses across environmental conditions was qualitatively similar irrespective of whether participants counted or ignored task stimuli. Nonetheless, sequential interaction contrasts revealed that counting accentuated the difference between standard and deviant responses in the Lab condition relative to the Field condition (*p*_adj_ =.028). A similar effect was noted for the Campus relative to the Field condition, on account of the increased difference between stimulus responses when participants counted deviants (*p*_adj_ =.038).

### P3 time-window

Type II Wald tests revealed a significant three-way interaction between stimulus, task, and environment for the P3 model (χ^2^ (2) = 33.40, *p* < .001; see Table 1 for full decomposition). This interaction is resolved in Figure 4 (see Table 2 for the full model summary).

Pairwise comparisons of the difference in responses to standards and deviants were significant across all factorial combinations of Task and Environment (all *p*_adj_ <.038), with the exception of the Ignore task in the Campus condition (*p*_adj_ =.588). Sequential interaction contrasts revealed that the difference between stimulus responses became progressively smaller from Lab, to Field, to Campus conditions in both task manipulations (all *p*_adj_ <.001). Counting served to increase the difference between stimulus responses (i.e. amplify the magnitude of the P3 effect) in each environmental condition (all *p*_adj_ <.001).

In contrast to our analysis of the MMN time-window, differences between the Count and Ignore tasks in the P3 time-window were primarily driven by modulations of deviant tone processing. Deviant stimuli elicited significantly stronger positivities in both the Lab and Field conditions when being counted as opposed to being ignored (all *p*_adj_ <.001). Counted and ignored deviants did not significantly differ from one another in the Campus condition (*p*_adj_ = .999).

## Discussion

The present study addressed the extent to which attentional processing is influenced by the situational context and its associated task demands. This involved comparing electrophysiological correlates of attentional processing between a traditional lab-based setting and two more naturalistic, dynamic scenarios involving walking and navigation. The manipulation of task instruction (i.e., to count or ignore oddball stimuli) across these three conditions enabled us to further compare the interaction of different levels of motor and environmental demands on top-down attentional engagement with task stimuli.

While participants were generally able to perform the oddball counting task with a high degree of accuracy across all three environmental conditions, linear mixed-effects regression analysis revealed that oddball counts were significantly less accurate in the Campus condition. This suggests the additional complexity entailed by this condition – both in terms of the more variable and unpredictable sensory stimuli it availed, and the more varied patterns of motor activity it solicited – were sufficiently distracting to degrade attentional focus on this simple cognitive task. By contrast, the additional motor demands and sensory complexity induced by walking around a sports field did not significantly affect task performance compared to sitting in a quiet room.

By contrast, the main findings from our ERP analyses revealed substantial differences in both the MMN and P3 components under ‘real-world’ versus laboratory conditions. This finding suggests that more ecologically-valid conditions involving motor co-ordination may modulate attentional processing despite ostensibly good behavioral task performance. Differences in ERPs between the Field and the Campus condition were subtle, which is perhaps surprising given the superior behavioural performance and relative simplicity of the Field condition. Differences between these conditions were limited to the Count task, and were more pronounced for the P3 response than the MMN.

With respect to previous studies of MMN and P3 components, findings from our Lab condition confirm the validity of the present results ^17,53,54^. Specifically, the pre-dominant distribution of activity across fronto-central and centro-parietal regions of the scalp, respectively, was in accordance with findings from dual-task walking studies ^55,56^, as well as basic studies of the MMN ^14,57^ and P3 ^58^.

Increasing evidence from previous laboratory as well as real-world studies point to the fact that attention on a given task depends on aspects such as task-difficulty, concurrent motor demands, as well as environmental conditions ^27^. However, especially in real-world scenarios findings are rare and partly controversial. The present findings tap into this gap – for example – by identifying distinctions in the effects of different real-world scenarios on attentional resources allocated to cognitive task-performance, by varying cognitive task difficulty. In contrast, previous studies with similar cognitive tasks reported no differences in P3 amplitude between different real-world scenarios ^3^. This apparent discrepancy can be explained by the varied characteristics of the real-world scenarios included within these studies. While simple conditions are comparable to the present sports field condition (e.g., walking on a circuit on the lawn in front of the university), more complex scenarios differ significantly from walking across the campus (e.g., obstacle course on an empty area or corridor).

Together with previous findings, we suggest that commonly-encountered everyday situations (e.g., wayfinding, interacting with other people, etc.) may place attentional resources under higher demand than physically-demanding situations (e.g., obstacle course) within a controlled, unchanging environment. In accordance with previous real-world EEG findings ^23,25,27^, evidence from laboratory studies, which used similar cognitive tasks, also identified no significant effects of increased motor demands (e.g., slow walking vs. fast walking on a treadmill) ^20^. However, studies with more difficult cognitive tasks (higher number of target-stimuli and increased complexity) report differences in amplitude, latency and topography of ERP components with increasing motor demands ^21^. It follows that the amount of attention deployed during a given cognitive task is influenced by the complexity of real-world situations, but more importantly, that the influence of environmental complexity is also modulated by cognitive task difficulty.

In contrast to previous studies, the present findings revealed significant interactions between stimulus, task, and condition for both ERP time-windows ^3^. The difference between standard and deviant stimulus responses diminished from lab to ‘real-world’ conditions. For the MMN, this effect was predominantly driven by a reduction in the positivity evoked by standard stimuli while walking; deviants, by contrast, evoked similar voltage levels irrespective of environmental or task condition. Analogously, standard stimuli evoked less-negative potentials in the P3 time-window during the walking conditions compared to the lab.

This pattern of results is broadly consistent with a predictive coding-style account of perceptual learning ^59,60^ and its attentional modulation via the mechanism of synaptic gain-control ^61,62^. Under the assumption that frequent, regular presentations of the standard tone enable the brain to construct a precise model of stimulus features (e.g., pitch, duration), the sensory signals generated by the repeated stimuli are efficiently cancelled-out (or ‘silenced’) by descending model predictions ^16,63^. The present findings suggest this ability may be reduced when moving through (and actively engaging with) more complex, dynamic environments. A possible explanation for this is that the additional, competing demands imposed on attention by more naturalistic scenarios may limit the brain’s capacity to accurately and precisely model recurrent stimuli, such that they are subject to more elaborate cortical processing than they would ordinarily be afforded under laboratory conditions. The functional upshot of this account is that standard stimuli may essentially be treated more like deviants when occasioned in the context of more complex, variable, and uncertain conditions than those typically encountered in the lab.

Currently, we do not fully understand the extent to which environmental stimulation and movement-related information recruit attention resources. However, based on the findings presented here and in other recently published naturalistic studies ^3,27^ it is becoming increasingly evident that factors such as gross motor activity (as opposed to sitting or standing), cognitive task difficulty, and the degree of experimental control over ambient conditions (i.e. laboratory vs. real-world settings), seem to be highly relevant determinants of attention and cognitive resource allocation. Such results highlight potential limitations in the generalizability of standard laboratory experiments to the more complex, dynamic conditions encountered in everyday life. However, it is also important to note some of the limitations and ongoing challenges faced by real-world designs, which we consider below.

The present study focused on EEG signals in order to avoid potential complications or confounds introduced by encumbering the participant with multiple measurement devices. One consequence of this decision was that we were unable to augment our analysis of cognitive task performance with information concerning gait parameters. Based on previous findings that identified a maintenance of cognitive performance in dual-task walking conditions by reducing walking speed ^64^, it can only be speculated that gait parameters in the present study may have changed with increased complexity of the situation (across the campus vs. around the sports field) and thereby, the amount of attention focused on the cognitive task could be maintained; however, this must remain a question for future research.

Moreover, while advancing technological innovations are revolutionizing the study of cognition under ecologically-valid conditions, mobile EEG designs still face a number of challenges. In addition to the concern that performing tasks and activities while wearing/carrying recording devices may perturb normal functioning, the amount and quality of the data collected under such conditions is often reduced compared to standard laboratory conditions. Although we attempted to mitigate this potential issue through the application of sophisticated artifact rejection techniques, epoch rejection rates varied considerably amongst individuals. The problem of noisy data collection might also have been compounded by the measurement imprecision introduced by the delayed (and in some cases, repeated) transmission of event markers. While we took care to correct these errors without introducing bias (i.e., applying a uniform adjustment designed to ameliorate event marker lag across all conditions prior to statistical analysis), the additional variance introduced into the data may have limited our sensitivity to fine-grained differences between conditions.

Despite these cautionary remarks, the rapid development of mobile EEG systems and software solutions suggests that many of the challenges mentioned above can be overcome in future studies. Methodological innovations such as the development of ear-EEG systems promise to enable the acquisition of high-quality data while participants move about freely ^65,66^. Sophisticated algorithms for capturing and characterizing gait-related artefacts are also being developed (see, e.g., ^67^). Such advancements will prove critical for enhancing the quality and robustness of mobile EEG data collected under complex and variable environmental conditions.

Existing findings from naturalistic EEG studies provide valuable insights concerning the nature of well-established markers of cognitive processing such as the MMN and P3 components. The present results align well with recently published mobile EEG studies and fill an important gap in this literature about the effects of environmental complexity, motor demands, and cognitive task load on attention within naturalistic scenarios. Furthermore, our findings highlight the necessity of extending future cognitive research programs beyond the confines of the laboratory. Ideally, such studies will combine complementary measurement devices as they become increasingly portable (e.g., ear-EEG & eye-tracking), and address a greater variety of everyday situations.

## Acknowledgements

AWC and CAH are supported by Australian Government Research Training Program (RTP) scholarships. IBS acknowledges the support of an Australian Research Council Future Fellowship (FT160100437). This research was funded by the University of South Australia under the Research Themes Investment Scheme. We would also like to thank the undergraduate students in course BEHL 3021 (“Cognitive Neuroscience”) at the University of South Australia who participated in this study and helped with data collection as part of their class project.

