## Supplemental material for "EEG and behavioral correlates of attentional processing while walking and navigating naturalistic environments"

Montage of ERP waveforms for each condition

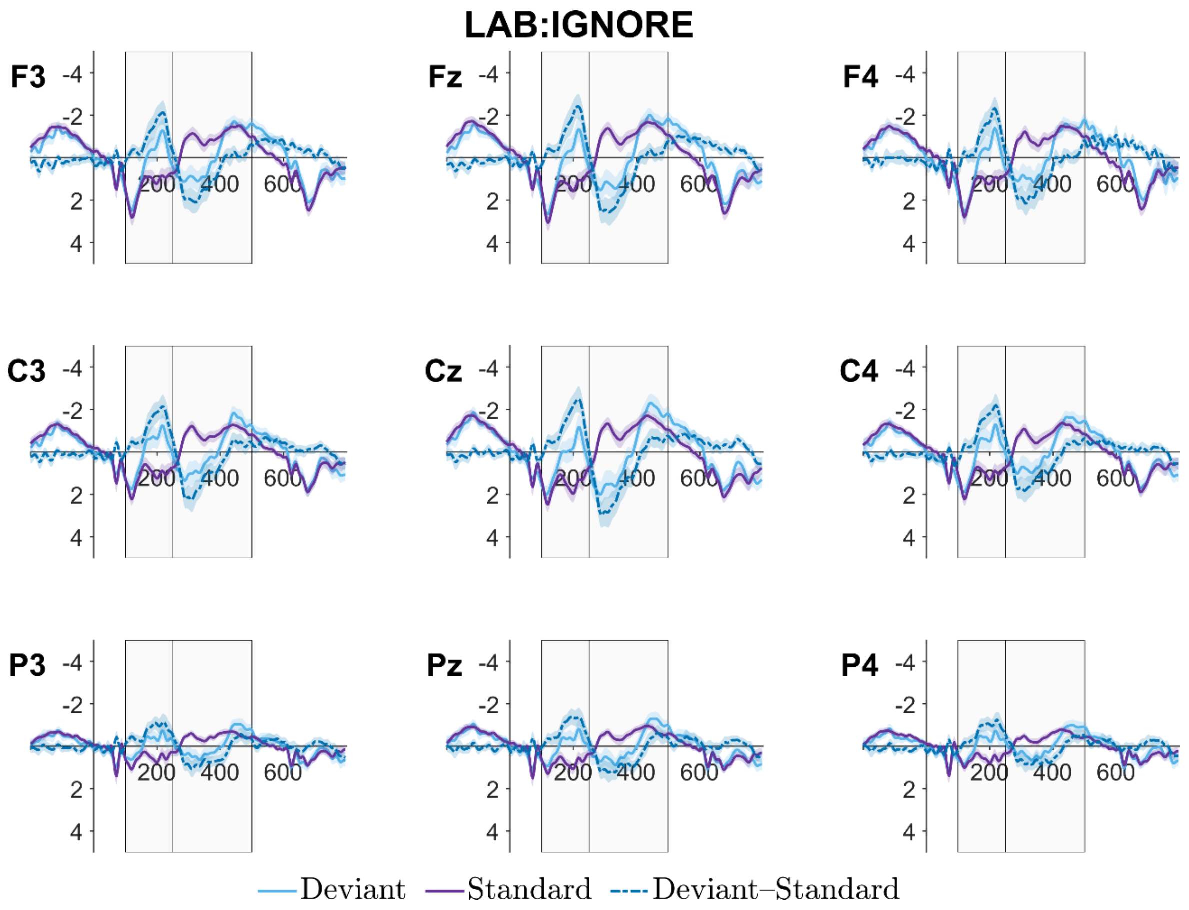


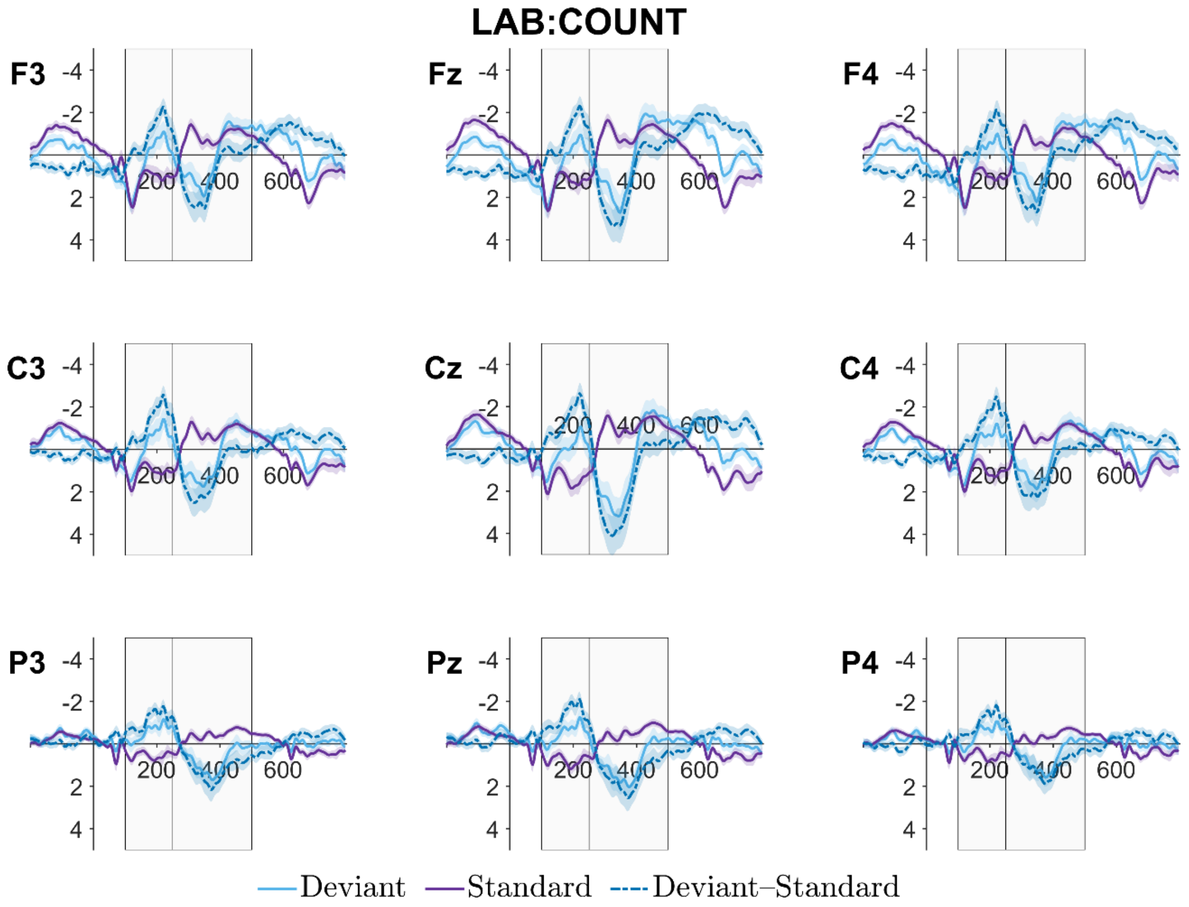

### FIELD:IGNORE

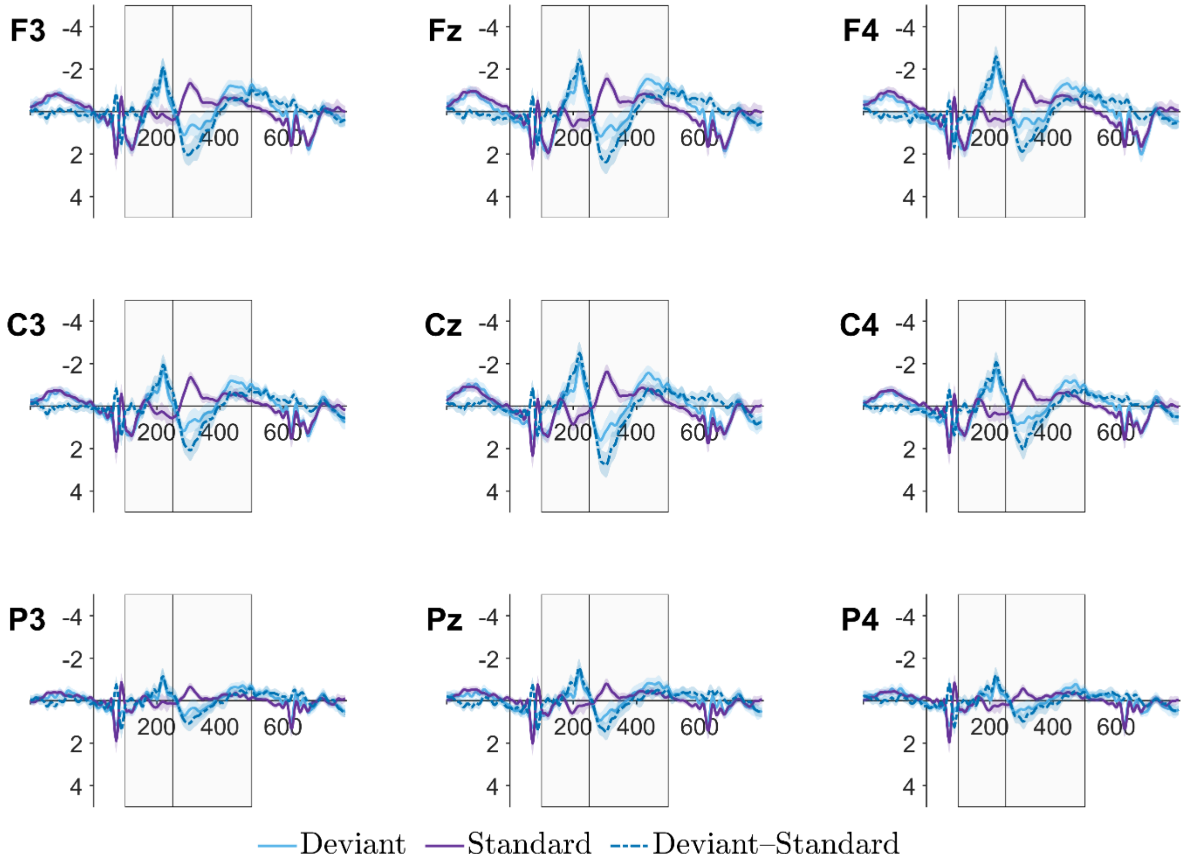


### FIELD:COUNT

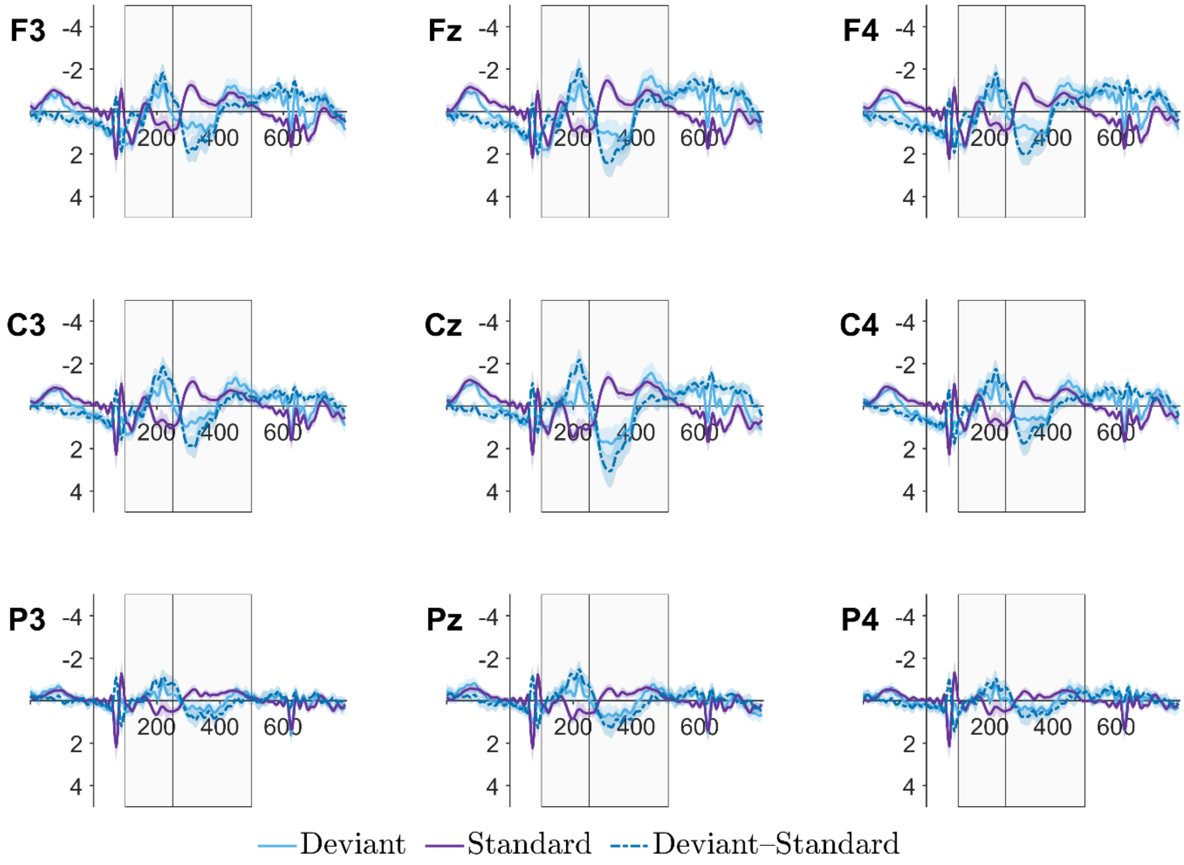

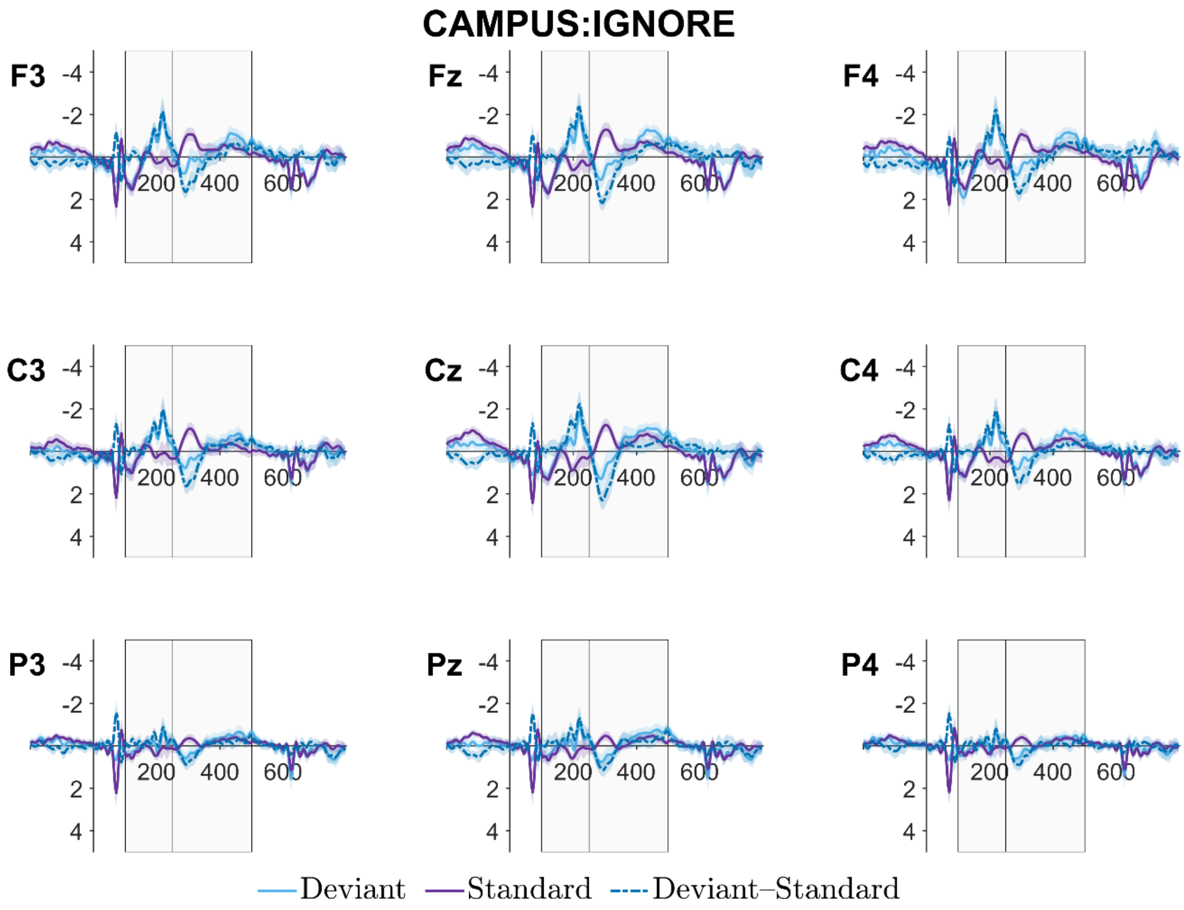
CAMPUS:IGNORE
F3
Fz
F4
C3
Cz
C4
P3
Pz
P4
-4
-2
2
4
200
400
600
Deviant
Standard
Deviant–Standard

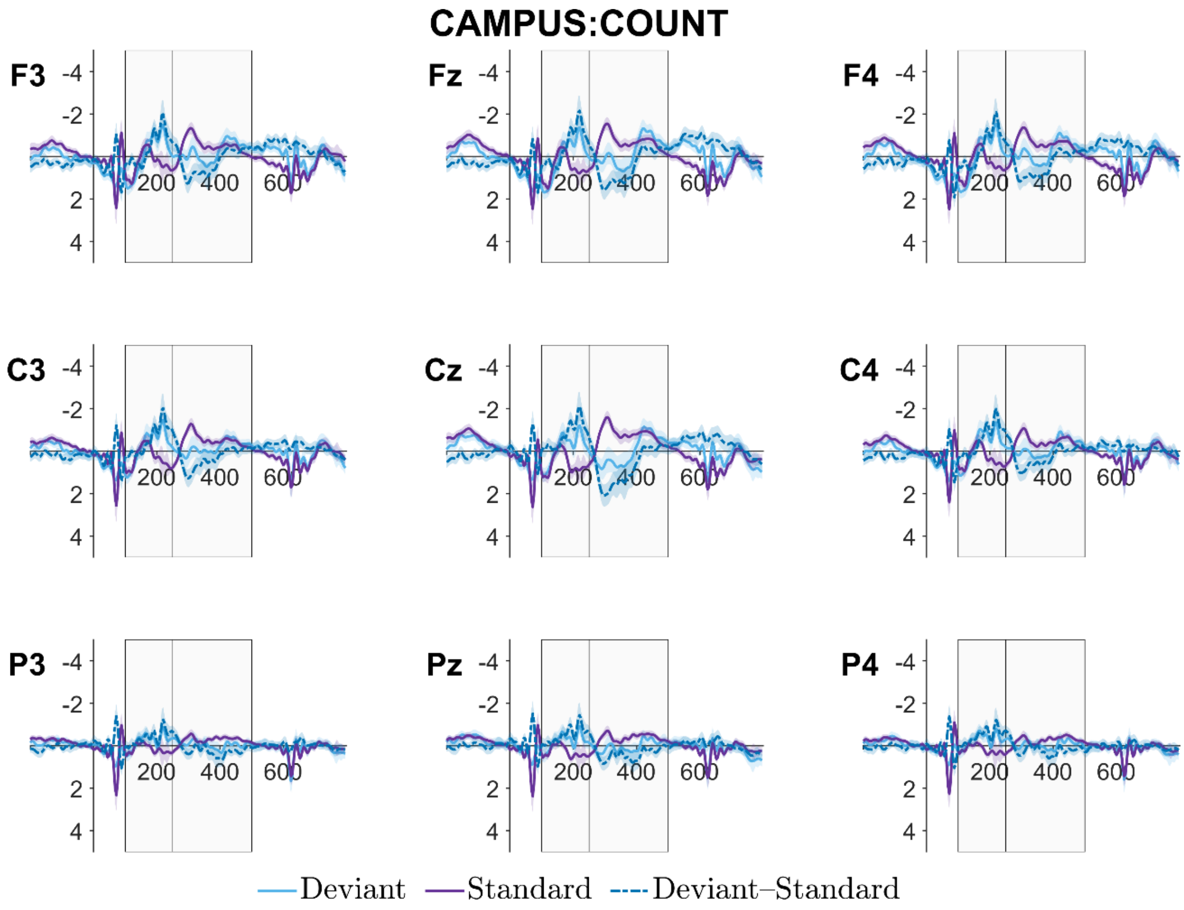
CAMPUS:COUNT
F3
Fz
F4
C3
Cz
C4
P3
Pz
P4
-4
-2
2
4
200
400
600
Deviant
Standard
Deviant–Standard

### Linear mixed-effects models of ERP time-windows factorised by midline channel location

The following model summaries and visualisations recapitulate the models reported in the main text using data from 4 midline channels only (Fz, Cz, Pz, Oz). Channel identity is included as a fixed (rather than random) effect and allowed to interact with other factorial variables.

| | **MMN time-window** | | | | **P3 time-window** | | | |
|---|---|---|---|---|---|---|---|---|
| *Predictors* | *Estimate* | *S.E.* | *t-stat* | *p-value* | *Estimate* | *S.E.* | *t-stat* | *p-value* |
| (Intercept) | 0.28 | 0.07 | 3.98 | **<0.001** | -0.23 | 0.06 | -3.56 | **0.001** |
| stimulus[S.Standard] | 0.28 | 0.01 | 30.11 | **<0.001** | -0.26 | 0.04 | -7.17 | **<0.001** |
| task[S.Ignore] | 0.00 | 0.01 | 0.13 | 0.898 | -0.07 | 0.01 | -9.38 | **<0.001** |
| environ[S.Lab] | 0.19 | 0.05 | 3.65 | **0.001** | -0.04 | 0.05 | -0.77 | 0.448 |
| environ[S.Field] | -0.10 | 0.03 | -3.04 | **0.004** | 0.04 | 0.03 | 1.18 | 0.246 |
| channel_id[S.Cz] | 0.12 | 0.02 | 7.41 | **<0.001** | -0.15 | 0.04 | -4.30 | **<0.001** |
| channel_id[S.Fz] | 0.20 | 0.02 | 12.31 | **<0.001** | -0.20 | 0.04 | -5.04 | **<0.001** |
| channel_id[S.Oz] | -0.17 | 0.02 | -10.40 | **<0.001** | 0.22 | 0.05 | 4.08 | **<0.001** |
| prestim.s | -1.39 | 0.01 | -200.96 | **<0.001** | -1.30 | 0.01 | -244.75 | **<0.001** |
| stimulus[S.Standard] : task[S.Ignore] | -0.01 | 0.01 | -1.03 | 0.301 | 0.07 | 0.01 | 9.78 | **<0.001** |
| stimulus[S.Standard] : environ[S.Lab] | 0.14 | 0.01 | 10.39 | **<0.001** | -0.17 | 0.01 | -16.80 | **<0.001** |
| stimulus[S.Standard] : environ[S.Field] | -0.05 | 0.01 | -3.53 | **<0.001** | 0.05 | 0.01 | 5.24 | **<0.001** |
| task[S.Ignore] : environ[S.Lab] | 0.04 | 0.01 | 3.23 | **0.001** | -0.10 | 0.01 | -9.42 | **<0.001** |
| task[S.Ignore] : environ[S.Field] | -0.06 | 0.01 | -4.89 | **<0.001** | 0.01 | 0.01 | 0.89 | 0.375 |
| stimulus[S.Standard] : channel_id[S.Cz] | 0.14 | 0.02 | 9.08 | **<0.001** | -0.17 | 0.01 | -13.91 | **<0.001** |
| stimulus[S.Standard] : channel_id[S.Fz] | 0.02 | 0.02 | 1.24 | 0.214 | -0.09 | 0.01 | -7.62 | **<0.001** |
| stimulus[S.Standard] : channel_id[S.Oz] | -0.21 | 0.02 | -12.42 | **<0.001** | 0.23 | 0.01 | 17.82 | **<0.001** |

| | | | | | | | | |
|---|---|---|---|---|---|---|---|---|
| task[S.Ignore] : channel_id[S.Cz] | -0.02 | 0.02 | -1.35 | 0.177 | -0.04 | 0.01 | -3.13 | **0.002** |
| task[S.Ignore] : channel_id[S.Fz] | -0.03 | 0.02 | -2.13 | **0.034** | 0.01 | 0.01 | 0.73 | 0.468 |
| task[S.Ignore] : channel_id[S.Oz] | 0.00 | 0.02 | 0.22 | 0.825 | 0.06 | 0.01 | 5.08 | **<0.001** |
| environ[S.Lab] : channel_id[S.Cz] | 0.14 | 0.02 | 6.31 | **<0.001** | -0.03 | 0.02 | -1.74 | 0.081 |
| environ[S.Field] : channel_id[S.Cz] | -0.07 | 0.02 | -3.14 | **0.002** | 0.03 | 0.02 | 1.75 | 0.080 |
| environ[S.Lab] : channel_id[S.Fz] | 0.17 | 0.02 | 7.52 | **<0.001** | -0.06 | 0.02 | -3.59 | **<0.001** |
| environ[S.Field] : channel_id[S.Fz] | -0.06 | 0.02 | -2.73 | **0.006** | 0.02 | 0.02 | 1.42 | 0.156 |
| environ[S.Lab] : channel_id[S.Oz] | -0.19 | 0.02 | -8.29 | **<0.001** | 0.02 | 0.02 | 1.05 | 0.292 |
| environ[S.Field] : channel_id[S.Oz] | 0.09 | 0.02 | 3.79 | **<0.001** | -0.01 | 0.02 | -0.49 | 0.623 |
| stimulus[S.Standard] : task[S.Ignore] : environ[S.Lab] | -0.01 | 0.01 | -0.79 | 0.429 | 0.04 | 0.01 | 3.84 | **<0.001** |
| stimulus[S.Standard] : task[S.Ignore] : environ[S.Field] | 0.03 | 0.01 | 1.93 | 0.054 | -0.01 | 0.01 | -1.38 | 0.168 |
| stimulus[S.Standard] : task[S.Ignore] : channel_id[S.Cz] | 0.01 | 0.02 | 0.35 | 0.724 | 0.05 | 0.01 | 3.84 | **<0.001** |
| stimulus[S.Standard] : task[S.Ignore] : channel_id[S.Fz] | 0.09 | 0.02 | 5.66 | **<0.001** | -0.03 | 0.01 | -2.39 | **0.017** |
| stimulus[S.Standard] : task[S.Ignore] : channel_id[S.Oz] | -0.03 | 0.02 | -1.59 | 0.111 | -0.05 | 0.01 | -3.99 | **<0.001** |
| stimulus[S.Standard] : environ[S.Lab] : channel_id[S.Cz] | 0.02 | 0.02 | 0.96 | 0.339 | -0.05 | 0.02 | -3.14 | **0.002** |
| stimulus[S.Standard] : environ[S.Field] : channel_id[S.Cz] | -0.00 | 0.02 | -0.10 | 0.921 | 0.01 | 0.02 | 0.33 | 0.738 |

| | | | | | | | | |
|---|---|---|---|---|---|---|---|---|
| stimulus[S.Standard] : environ[S.Lab] : channel_id[S.Fz] | -0.03 | 0.02 | -1.39 | 0.164 | -0.03 | 0.02 | -1.50 | 0.134 |
| stimulus[S.Standard] : environ[S.Field] : channel_id[S.Fz] | 0.01 | 0.02 | 0.50 | 0.616 | -0.00 | 0.02 | -0.14 | 0.885 |
| stimulus[S.Standard] : environ[S.Lab] : channel_id[S.Oz] | -0.05 | 0.02 | -2.29 | **0.022** | 0.11 | 0.02 | 6.23 | **<0.001** |
| stimulus[S.Standard] : environ[S.Field] : channel_id[S.Oz] | 0.02 | 0.02 | 0.75 | 0.452 | -0.03 | 0.02 | -1.44 | 0.151 |
| task[S.Ignore] : environ[S.Lab] : channel_id[S.Cz] | 0.02 | 0.02 | 0.74 | 0.462 | 0.01 | 0.02 | 0.32 | 0.752 |
| task[S.Ignore] : environ[S.Field] : channel_id[S.Cz] | -0.03 | 0.02 | -1.53 | 0.127 | -0.02 | 0.02 | -1.00 | 0.318 |
| task[S.Ignore] : environ[S.Lab] : channel_id[S.Fz] | -0.03 | 0.02 | -1.50 | 0.134 | 0.05 | 0.02 | 2.92 | **0.003** |
| task[S.Ignore] : environ[S.Field] : channel_id[S.Fz] | 0.03 | 0.02 | 1.39 | 0.165 | -0.05 | 0.02 | -3.10 | **0.002** |
| task[S.Ignore] : environ[S.Lab] : channel_id[S.Oz] | -0.01 | 0.02 | -0.27 | 0.786 | -0.01 | 0.02 | -0.58 | 0.560 |
| task[S.Ignore] : environ[S.Field] : channel_id[S.Oz] | 0.00 | 0.02 | 0.09 | 0.929 | 0.04 | 0.02 | 2.25 | **0.025** |
| stimulus[S.Standard] : task[S.Ignore] : environ[S.Lab] : channel_id[S.Cz] | 0.01 | 0.02 | 0.61 | 0.542 | -0.01 | 0.02 | -0.64 | 0.523 |
| stimulus[S.Standard] : task[S.Ignore] : environ[S.Field] : channel_id[S.Cz] | -0.01 | 0.02 | -0.57 | 0.567 | 0.02 | 0.02 | 1.14 | 0.254 |
| stimulus[S.Standard] : task[S.Ignore] : | 0.03 | 0.02 | 1.39 | 0.166 | -0.04 | 0.02 | -2.31 | **0.021** |

| | | | | | | | | |
|---|---|---|---|---|---|---|---|---|
| environ[S.Lab] : channel_id[S.Fz] | | | | | | | | |
| stimulus[S.Standard] : task[S.Ignore] : environ[S.Field] : channel_id[S.Fz] | -0.02 | 0.02 | -0.78 | 0.433 | 0.04 | 0.02 | 2.40 | **0.017** |
| stimulus[S.Standard] : task[S.Ignore] : environ[S.Lab] : channel_id[S.Oz] | -0.01 | 0.02 | -0.59 | 0.554 | 0.02 | 0.02 | 0.89 | 0.374 |
| stimulus[S.Standard] : task[S.Ignore] : environ[S.Field] : channel_id[S.Oz] | 0.04 | 0.02 | 1.49 | 0.137 | -0.04 | 0.02 | -2.43 | **0.015** |
| **Random Effects** | | | | | | | | |
| $\sigma^2$ | 31.70 | | | | 18.60 | | | |
| $\tau_{00}$ | 0.17 subject_id | | | | 0.15 subject_id | | | |
| $\tau_{11}$ | 0.09 subject_id.environ[S.Lab] | | | | 0.04 subject_id.stimulus[S.Standard] | | | |
| | 0.03 subject_id.environ[S.Field] | | | | 0.09 subject_id.environ[S.Lab] | | | |
| | | | | | 0.04 subject_id.environ[S.Field] | | | |
| | | | | | 0.04 subject_id.channel_id[S.Cz] | | | |
| | | | | | 0.05 subject_id.channel_id[S.Fz] | | | |
| | | | | | 0.10 subject_id.channel_id[S.Oz] | | | |
| $\rho_{01}$ | 0.05 subject_id.environ[S.Lab] | | | | -0.19 subject_id.stimulus[S.Standard] | | | |
| | 0.12 subject_id.environ[S.Field] | | | | 0.20 subject_id.environ[S.Lab] | | | |
| | | | | | 0.03 subject_id.environ[S.Field] | | | |
| | | | | | 0.70 subject_id.channel_id[S.Cz] | | | |
| | | | | | 0.74 subject_id.channel_id[S.Fz] | | | |
| | | | | | -0.83 subject_id.channel_id[S.Oz] | | | |
| N | 36 subject_id | | | | 36 subject_id | | | |
| Observations | 572291 | | | | 572291 | | | |

*Note*. Factor variables sum-to-zero contrast-coded. *Estimate* = beta coefficient, *S.E.* = standard error, *stimulus* = oddball task stimulus (Deviant, Standard), *task* = cognitive task condition (Count, Ignore), *environ* = environmental context (Lab, Field, Campus), *channel_id* = electrode channel location (Fz, Cz, Pz, Oz), *prestim.s* = prestimulus EEG voltage (scaled), $\sigma^2$ = residual variance, $\tau_{00}$ = random intercept variance, $\tau_{11}$ = random slope variance, $\rho_{01}$ = random effect correlation coefficient, *N* = number of levels per grouping variable.

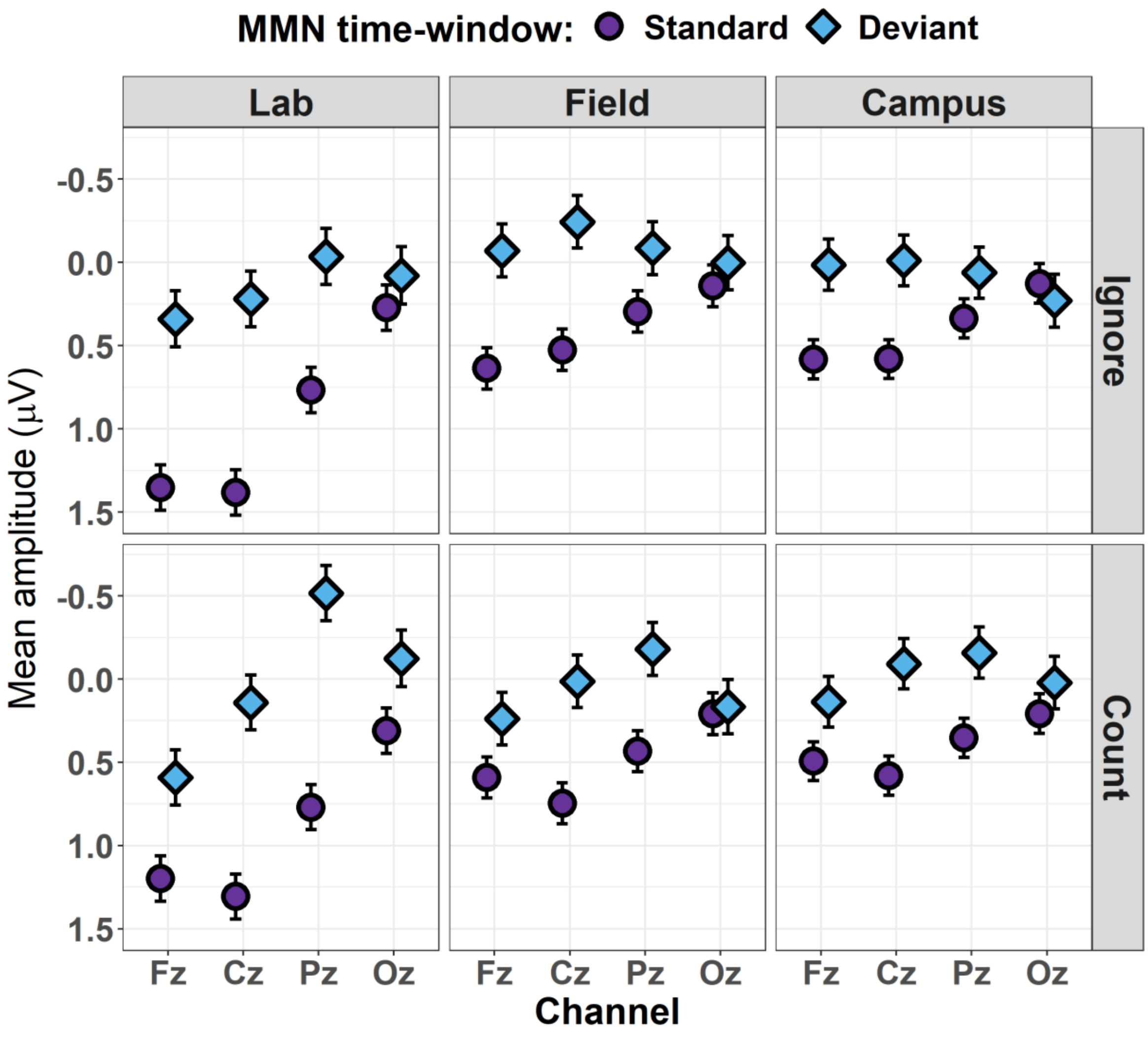


Visualisation of estimated marginal mean voltages for the MMN time-window (100-250 ms) factorized by stimulus (Standard, Deviant), task (Ignore, Count), environment (Lab, Field, Campus), and channel location (Fz, Cz, Pz, Oz). Error bars indicate 84% confidence intervals.

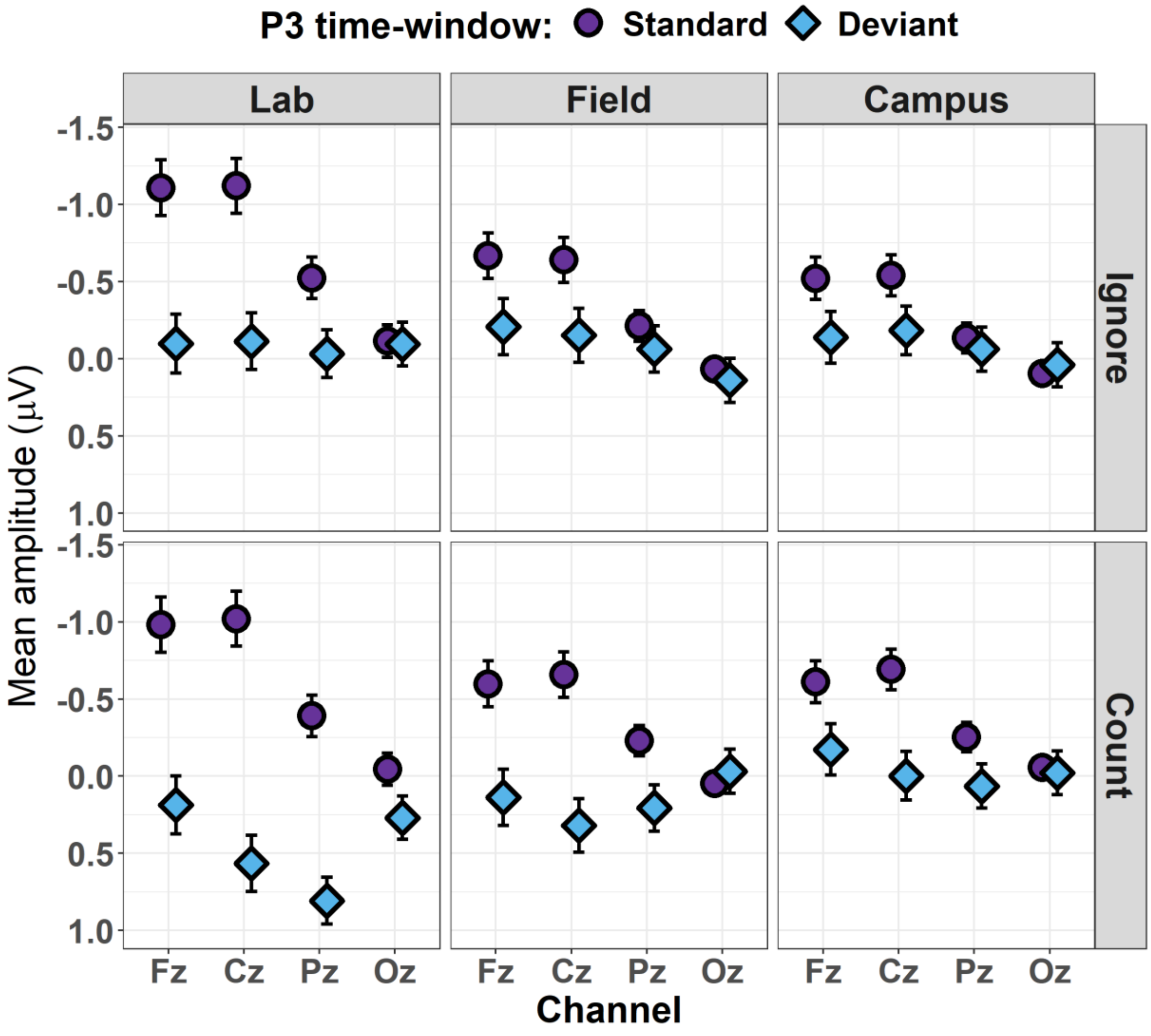


Visualisation of estimated marginal mean voltages for the P3 time-window (250-500 ms) factorized by stimulus (Standard, Deviant), task (Ignore, Count), environment (Lab, Field, Campus), and channel location (Fz, Cz, Pz, Oz). Error bars indicate 84% confidence intervals.
